# Unsupervised detection of cell-assembly sequences with edit similarity score

**DOI:** 10.1101/202655

**Authors:** Keita Watanabe, Tatsuya Haga, David R Euston, Masami Tatsuno, Tomoki Fukai

## Abstract

Cell assembly is a hypothetical functional unit of information processing in the brain. While technologies for recording large-scale neural activity have been advanced, mathematical methods to analyze sequential activity patterns of cell-assembly are severely limited. Here, we propose a method to extract cell-assembly sequences repeated at multiple time scales and various precisions from irregular neural population activity. The key technology is to combine “edit similarity” in computer science with machine-learning clustering algorithms, where the former defines a “distance” between two strings as the minimal number of operations required to transform one string to the other. Our method requires no external references for pattern detection, and is tolerant of spike timing jitters and length irregularity in assembly sequences. These virtues enabled simultaneous automatic detections of hippocampal place-cell sequences during locomotion and their time-compressed replays during resting states. Furthermore, our method revealed previously undetected cell-assembly structure in the rat prefrontal cortex during goal-directed behavior. Thus, our method expands the horizon of cell-assembly analysis.

## INTRODUCTION

Uncovering neural codes is of fundamental importance in neuroscience. Several experimental results suggest that synchronous or sequential firing of cortical neurons play active roles in primates (Abeles et al., 1993; Hatsopoulos et al., 1998; Riehle et al., 1997; Steinmetz et al., 2000). In the rat somatosensory and auditory cortices, spontaneous and stimulus-evoked activities exhibit repeating sequences of neuronal firing (Luczak et al., 2009; 2007). In rodent hippocampus, place cells exhibit precisely-timed, repeating firing sequences representing the rats trajectory, subsections of which repeat during each theta cycle (Mehta et al., 2002; O’Keefe, 1976; Villette et al., 2015). These sequences are replayed at compressed temporal scales during awake immobile and sleep states (Buzsáki and Moser, 2013; Carr et al., 2011; Foster and Wilson, 2006; Lee and Wilson, 2002; Pfeiffer and Foster, 2013; Skaggs and McNaughton, 1996) presumably for memory consolidation (Girardeau et al., 2009; Jadhav et al., 2012). Similar replay events have also been observed in the rodent prefrontal cortex (Euston et al., 2007).

The rapid development of techniques for large-scale recording of neuronal activity provide fertile ground for the analysis of spike sequences. Calcium imaging enables simultaneous measurement of spike rates from hundreds to thousands of neurons (Deneux et al., 2016; Greenberg et al., 2008; Kerr et al., 2005; Mukamel et al., 2009; Ozden et al., 2008; Pnevmatikakis et al., 2016; Sasaki et al., 2007; Vogelstein et al., 2010), and imaging by voltage indicators may further overcome the poor temporal resolution in imaging (Emiliani et al., 2015; Grinvald and Petersen, 2015; Knöpfel et al., 2015). Extracellular recording of membrane potentials with multi-electrodes has also evolved, allowing access to spike trains from large numbers of neurons (Buzsáki, 2004; Buzsáki et al., 2015; Einevoll et al., 2012).

Despite this progress in experimental techniques, methods for analyzing the spatiotemporal structure of cell assemblies are still limited (Chen and Wilson, 2017). Template matching is a standard technique for the detection of repeated activity patterns (Abeles et al., 1993; Euston et al., 2007; Greenberg et al., 2008; Kerr et al., 2005; Luczak et al., 2007; 2009; Mukamel et al., 2009; Sasaki et al., 2008; Tatsuno et al., 2006; Vogelstein et al., 2010). However, the method requires reference events, such as sensory cues and motor responses, and is easily disrupted by biological noise, such as jitters in spike timing and variation in sequence length. On the other hand, only a few studies have attempted the blind detection of cell-assembly sequences without relying on reference events (Humphries, 2011; Picado-Muiño et al., 2013; Russo et al., 2017; Shimazaki et al., 2012), and such data analysis remains a challenge.

Here, we develop a novel method to detect self-similar firing patterns within cell assemblies by using the edit similarity score. Edit similarity is a metric originally introduced in computer science to measure the distance between arbitrary strings, and has been utilized for analyzing various types of sequences in network science and biology. Edit similarity measures matching between two sequences with flexible temporal alignment, which is an essential feature for detecting noisy spatiotemporal patterns embedded in neural activity. We extend the edit similarity score to a form applicable to neural activity data, and develop a clustering method for blind cell-assembly detection. We evaluated the performance of the method with artificial data, and found that our method is more robust against background noise than conventional clustering methods. Furthermore, we applied our method to experimental data recorded from the rat hippocampus (Mizuseki et al., 2009) and prefrontal cortex (Euston et al., 2007), and the algorithm detected several multi-cell sequences linked with behavior in an unsupervised manner. Robustness to noise and computational efficiency of our method will help the exhaustive search of repeated spatiotemporal patterns in large-scale neural data, which may lead to the elucidation of hidden neural codes.

## RESULTS

Our goal is to develop a method for detecting similar temporal patterns repeatedly occurring in neural population activity without relying on external sensory or behavioral events. Each of these patterns may represent a sequence of neural ensembles coincidently firing in irregularly repeated temporal windows of a certain length (Figure 1A). The detection of sequentially activated cell assemblies is difficult because repeated sequences are usually not exactly the same (Figure 1B): (i) temporal structure of the target sequence is shown; (ii) sequences may be contaminated by noisy spikes belonging to none of the cell assemblies; (iii) sequences may not always repeat a complete set of cells; (iv) repeated sequences of the member neurons may exhibit large jitters; (v) sequences can be expanded or compressed from one repetition to another; and (vi) different sequence may share the same neurons. Furthermore, some neurons may belong to multiple sequences. Such overlaps further complicate sequence detection.

**Figure 1:**
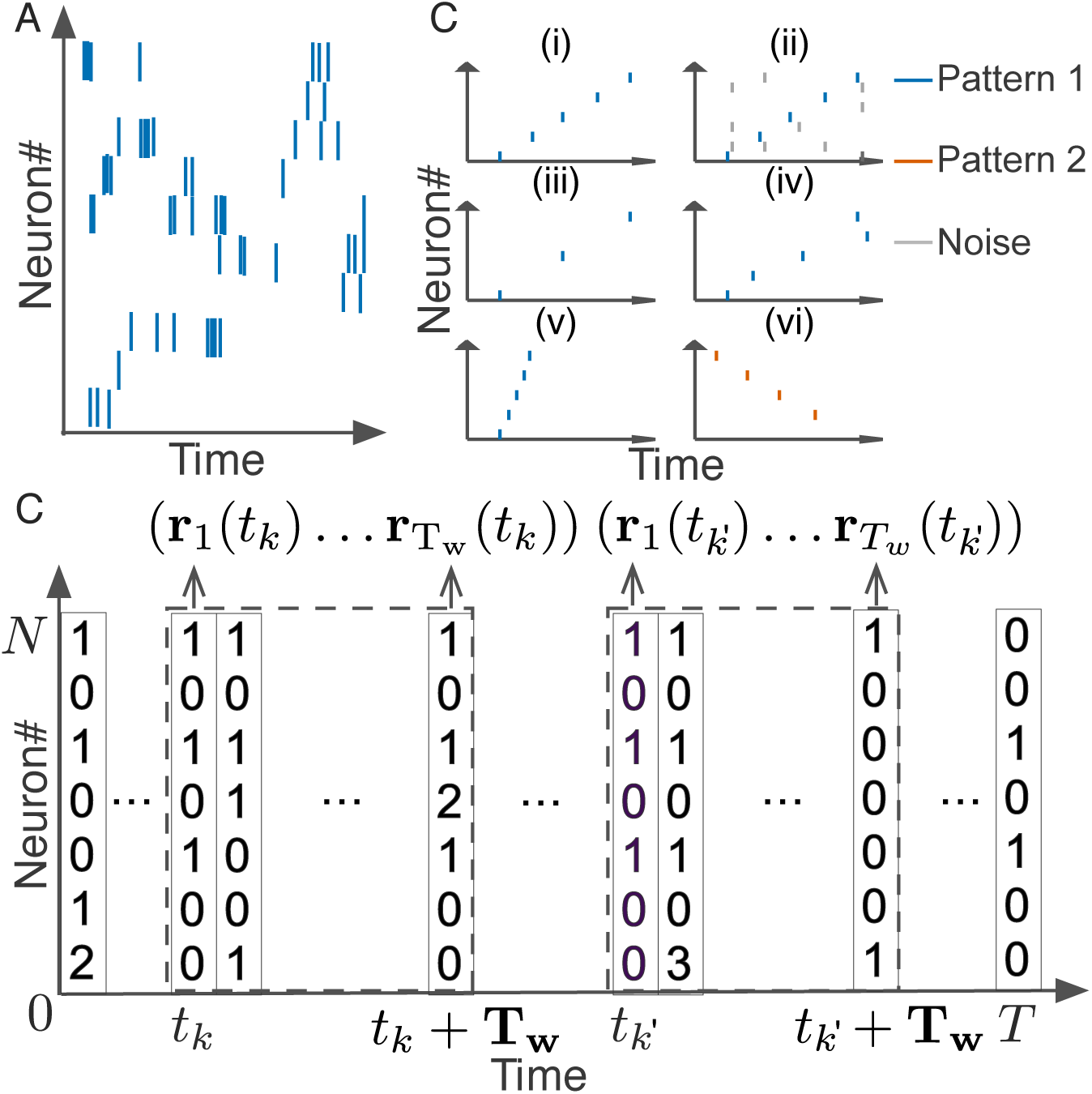
Detection of repetitive cell-assembly sequences. (A) A temporal pattern in a sliding time window is schematically illustrated. Such a pattern may contain neurons belonging to a cell assembly as well as non-member neurons. Member neurons may fire at different rates with different temporal precision. Similarity between cell assembly sequences will increase when they share more member neurons and when they fire in a more similar temporal order at similar firing rates with higher temporal precision. Note that each member neuron may appear multiple times at different temporal positions in a time window. (B) Difficulties in detecting repetitive cell-assembly sequences are schematically illustrated. Blue and red bars show spikes of member neurons, while gray bars represent noisy spikes of non-member neurons: (i) temporal structure of target sequence; (ii) contamination by spikes of non-member neurons; (iii) missing spikes of member neurons; (iv) jitters in spike timing of member neurons; (v) arbitrary scaling of sequence length; (vi) member overlaps between different sequences. (C) Sliding time windows W(t_k_) are divided into L bins with an identical size, where t_k_ refers to the start time of the k-th time window. Population rate vector consists of the spike counts of individual neurons in each bin.

### Extended edit similarity score between cell-assembly sequences

To overcome these difficulties, we developed a novel method for robust sequence detection based on the edit similarity score known in computer science (Levenshtein, 1966). Suppose that we evaluate similarity between two strings of genes, “ATCGTAC” and “ATGTTAT”. We may naively count the number of coincident bases at the corresponding positions in the two strings. In the above example, the first two bases “AT” coincide, so the similarity is two. However, if we count the maximal number of coincidences preserving the serial orders of bases but allowing the insertion of blanks “-”, we may compare “ATCGT-A-C” and “AT-GTTAT-” to obtain the maximal number of five (i.e., A, T, G, T and A coincide in this order). This explains the basic concept of edit similarity score.

We modified the Needleman-Wunsch (N-W) algorithm (Needleman and Wunsch, 1970) for scoring edit similarity such that it is applicable to neural data (see Experimental Procedures for details). Briefly, we segmented data with a sliding time window of width *T*_*w*_, and divided each data segment into *L* bins with size *b* (thus, *L* = *T*_*w*_/*b*). If we consider the activity pattern of the neural ensemble in each bin (i.e., the rate vector *r* of coincidently firing neurons in Figure 1C) as a “letter”, we obtain a string of letters in each data segment. Note that each neuron may fire multiple spikes in a bin, so each component of the activity vector represents the number of spikes generated by the corresponding neuron in the bin. The window size and bin size are determined from the temporal features of the neural data. In this study, the typical values of *T*_*w*_ ranged from 100ms to 250ms and those of *b* from 1 ms or 10 ms.

Our task is to find all data segments that contain similar activity vectors in the same temporal order (Figure 1C). In the previous comparison of two gene sequences, how to count the number of coincident letters (i.e., nucleotide bases) between the two sequences was naturally defined. However, the same scoring scheme is not applicable to neural activity data because neural population in vivo will hardly repeat exactly the same patterns due to various noise sources. In this study, we extended the edit similarity score using the inner product of activity vectors (Experimental Procedures).

Another important characteristic of our algorithm is an exponentially growing penalty. Edit similarity score is tolerant to various types of noise such as spike timing jitters, failure of firing, contamination of noisy spikes, and temporal variation in sequence length. However, even if their temporal orders are identical, the score should be reduced when two sequences involve significantly different time lags between the corresponding successive member neurons. To systematically reduce the score in such cases, we subtract a penalty term from the score whenever we inserted an extra blank (time bin) into one of the two sequences to increase the match between them. If continuous blanks are inserted, the net penalty grows exponentially as the length of inserted blanks is increased. The score is set equal to zero if the subtraction results in a negative value (Experimental Procedure).

Finally, the proposed method, in its original form, requires extensive computational resources when used on neural data. The major difficulty comes from the calculation of a similarity matrix that has a computational complexity of *O*(*M* ^2^), where M is the number of data segments and grows with the data length *T*. We accelerated the calculation of edit similarity drastically by employing an approximation algorithm based on Jaccard similarity (Cohen et al., 2001). For instance, this algorithm reduced the computation time by approximately 97% on the hippocampal data analyzed here. The details are explained in Experimental Procedures.

### Density- and community-based algorithms for sequence clustering

To find the data segments containing similar sequential activity patterns, we introduced a high dimensional metric space in which data segments form a cluster of neighboring data points. We can use the edit similarity score to define a metric among data points in this feature space (Experimental Procedures). While similar activity patterns give a dense cluster in the feature space, data segments containing no repeated patterns are scattered over the space as outliers. To remove these “noisy” components, we combined two different types of clustering algorithms. The first algorithm is called “OPTICS” and it finds dense clusters of data points (Ankerst et al., 1999). However, this algorithm cannot discriminate two clusters if they share a non-negligible number of data points. Such an example is shown in Figure 2A for an artificial dataset in which the algorithm identifies a single dense cluster and removes noisy components surrounding the cluster, but it does not separate the cluster into two parts.

**Figure 2:**
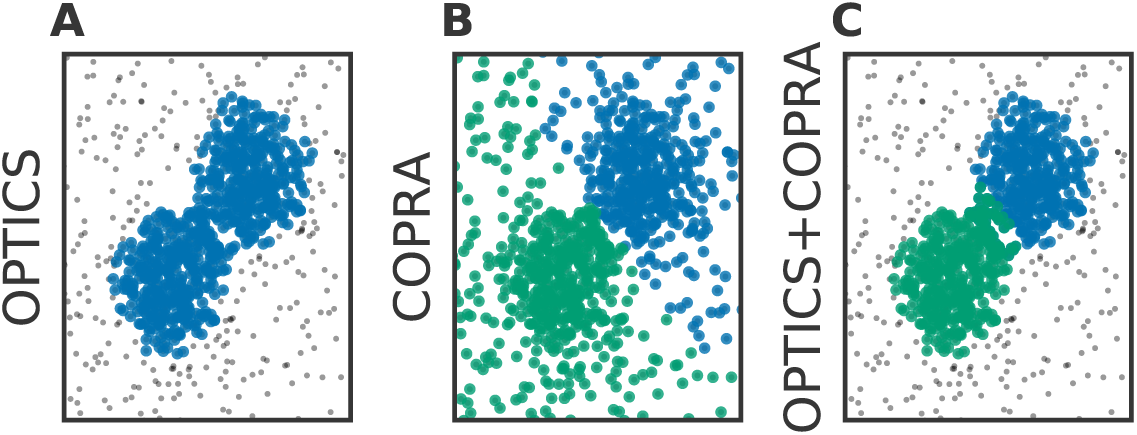
Performance comparison between different clustering algorithms. A density-based clustering algorithm (OPTICS) and a community detection algorithm (CORPA) were separately (A and B) and sequentially (C) applied to an identical dataset. Data points (blue, green) generated by two Gaussian distributions with different centers and an identical variance were mixed with uniformly distributed background data points (gray). (A) OPTICS could remove background noise but failed to discriminate the two clusters. (B) COPRA could separate these clusters but failed to remove background noise. (C) The combined application of OPTICS and COPRA successfully separated the two clusters and removed background noise.

The other algorithm “COPRA” performs clustering based on a community detection scheme (Gregory, 2010). In short, the data points connected with relatively short distances are distinguished from other data points as a community in the feature space. The algorithm identified two separate clusters in the same dataset as used above (Figure 2B). Each cluster, however, contained a considerable number of noisy data points: an outlier may be invited to a community if its distance from any member of the community is short enough. In this study, we sequentially applied OPTICS and CORPA to take advantage of each method. The combination of the two methods worked efficiently in most of the cases tested here (Figure 2C).

### Profiling cell-assembly sequences

We devised a method to identify the core temporal structure of the cell assembly sequences belonging to each cluster. Our method finds clusters of data segments within a neural ensemble that contain similar activity patterns, but these patterns in general exhibit large temporal variation. To find the core temporal cell-assembly structure for each cluster, called the “profile” in this study, we modified an iterative algorithm proposed previously (Barton and Sternberg, 1987). First, we choose two initial activity patterns from a cluster and construct a tentative profile from the sequence elements that commonly appear in the two patterns. Then, we sample another pattern that best matches this tentative profile of the cluster, and construct a new profile by taking the common elements between the tentative profile and the new sample. We repeat this updating procedure until the profile converges to a fixed pattern. For the applications reported in this study, updating 10 to 100 times yielded sufficiently good results. See Experimental Procedures for the details of our profiling procedure.

### Comparison between different methods on artificial data

We compared the performance of our method with that of PCA- (Lopes-dos-Santos et al., 2011; Peyrache et al., 2009) or ICA-based method (Lopes-dos-Santos et al., 2013) by using synthetic population activity data. We embedded 5 non-overlapping cell assemblies into background activity constructed using 100 simulated neurons firing independently at a rate of 2 [Hz]. Each cell assembly consisted of 20 synchronously firing neurons with ±10 milliseconds jitter and appeared 50 times at randomly determined positions within the data length of 300 seconds. The cell assemblies detected by our method are shown in Figure 3A. We evaluated the performance of each method in terms of F measure, which is widely used in the field of machine learning (c.f. (Artiles et al., 2007)~:

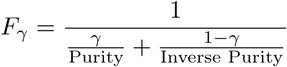

where *γ* is the mixing weight for Purity and Inverse Purity, Purity is a weighted average of the fractions of true members in detected clusters, and Inverse Purity is a weighted average of correctly classified portions of true clusters. If a classification is perfect, both Purity and Inverse Purity take the maximal value of unity. Other details of the evaluation method are described in **Experimental Procedures**.

**Figure 3:**
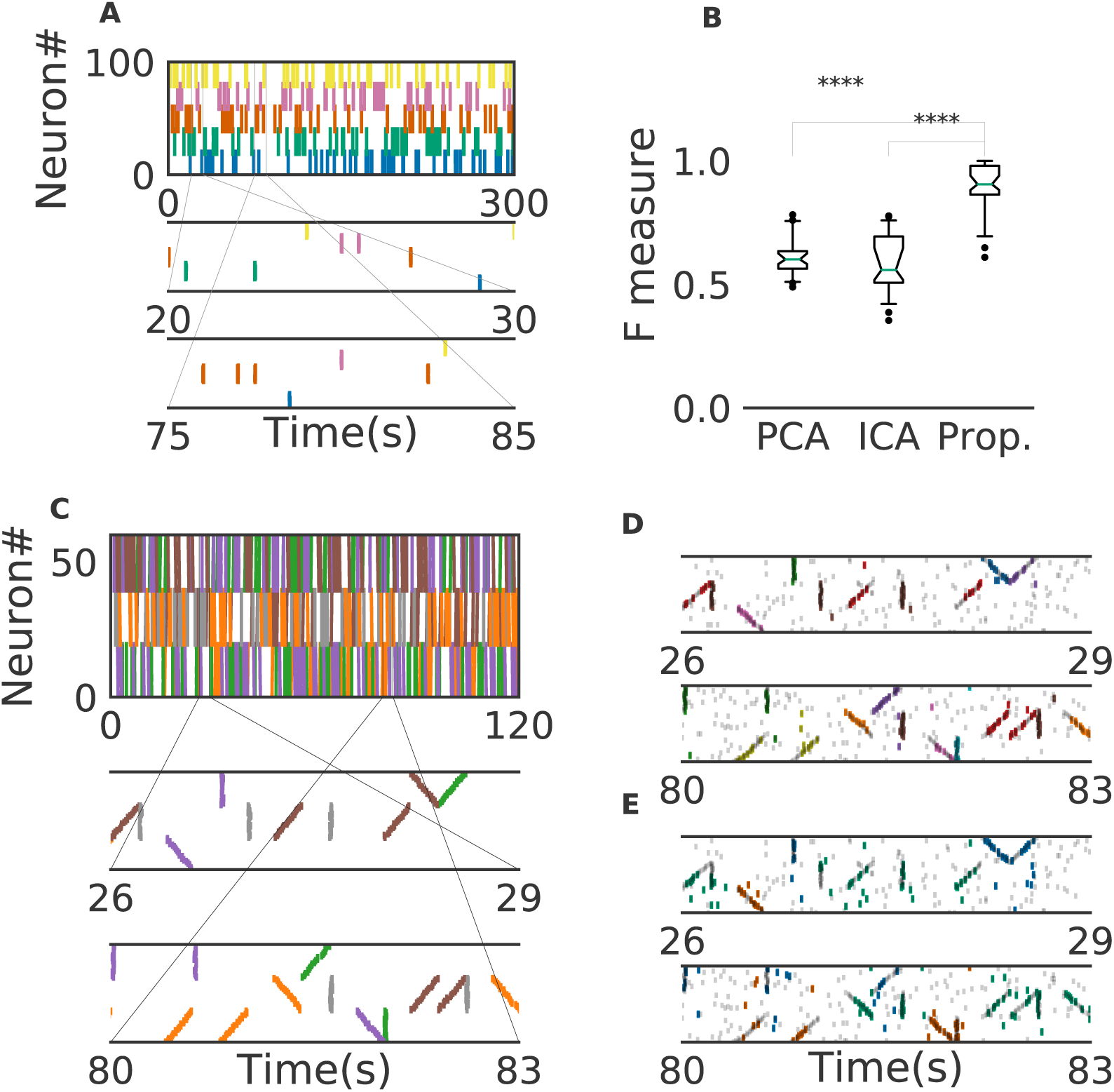
Comparison between different methods. (A) An example of the embedded artificial cell assemblies used for the comparison. In the raster plot, each dot is a spike. Each color indicates a cell assembly. For clarity, massively many noisy spikes are not shown. (B) F-score was significantly higher for the proposed method than for PCA-and ICA-based algorithms (p < 0.0001, Welch's t-test). The time window used was 200ms. (C) Nine artificial spike sequences embedded into noisy spike trains are shown. Noise spikes are not shown here. The time window was 200 ms. (D) Sequences detected by the proposed method from the artificial data shown in C are shown in two intervals together with all noisy spikes (grey). The nine embedded sequences were successfully detected. (E) Sequences detected from the same artificial data are shown for different values of parameters in our algorithm.

We generated 40 different artificial data by changing background activity. We then analyzed each data by three different methods and calculated *F*_1/2_ (the harmonic mean of Purity and Inverse Purity) for each trial. The resultant F measure was significantly larger in the proposal method (Mean ± Std 0.89±0.09) than in the PCA-based (0.61±0.07) and ICA-based (0.59±0.11) methods (Figure 3B). In fact, our method correctly detected all target cell assemblies.

We also investigated if our method is able to extract spike sequences in noisy artificial data. Sequential firing of three cell assemblies each consisting of 20 neurons was embedded into background activity at a rate of 1[Hz] in both forward, synchronous and reverse orders with ±10 milliseconds jitter(Figure 3C). Each sequences appeared 20 times. In Figures 3D and 3E, different values of parameters were tested to study the parameter dependence of our method, where criteria for timing jitters and clustering are stricter in Figure 3D (Experimental Procedure). In Figure 3D, our method detected the groups of cells which fire sequentially but in different orders as separate clusters (e.g., orange and red clusters). In contrast, the same group of cells firing in different orders was detected as a single cluster in Figure 3E. Namely, three separate groups of cells can fire with either an upward ramp, a downward ramp, or all simultaneously. Although some spikes are misidentified, the algorithm correctly identified each of the three clusters as was indicated by the colors. In summary, the investigation demonstrated the flexibility of the proposed method that would be useful for the investigation of memory reactivation; it can detect the forward and reverse replays separately or together depending on the researcher’s need.

### Place-cell firing sequences in the hippocampus

We now demonstrate that our method enables an automatic detection of firing sequences of rat hippocampal neurons during spatial exploration (Mizuseki et al., 2009). The data is available at the data sharing website of Collaborative Research in Computational Neuroscience (CRCNS.org., http://dx.doi.org/10.6080/K09G5JRZ). The data contains the activity of 108 neurons recorded from the CA1 region of the rat hippocampus during voluntary exploration of a linear maze. The total duration of recordings is 1928 seconds. See Experimental Procedures for the choices of parameter values.

Our method extracted 60 distinct clusters of data segments, each of which appeared repeatedly at a different location in the maze and in a specific movement direction (Figure 4A). We separated these clusters in terms of their observed locations. Twenty clusters appeared when the rat ran from the start to the goal (we call it the Go cluster), 20 clusters appeared during the opposite movement (called the Back cluster). There are five overlapping clusters between the two types (#6, #10, #21, #40, #48). Another 20 clusters mainly appeared during immobilility. The relationship between each cluster and a behavioral state indicates that our method successfully detected behaviorally relevant cell assemblies, which likely consist of hippocampal place cells. Some clusters (e.g., clusters #4 and #10) were detected during both locomotion and the resting state. These patterns presumably correspond to place-cell firing phase-locked to theta oscillation and their ripple-associated replays, respectively (Foster and Wilson, 2006; Nádasdy et al., 1999), and it is notable that our method automatically extracted these sequences in spite of the different time scale. Such reactivation was also observed in (Mizuseki et al., 2009). Figure 4B shows visualization of the feature space with t-SNE, which defines a mapping from high-dimensional data space to a low-dimensional space for visualization such that the spatial relationships between data points are optimally preserved (Maaten and Hinton, 2008). Figure 4C displays 4 sample profiles of activity patterns for the clusters detected (Experimental Procedures), where only the first 10 neurons are shown. Three examples (#4, #16, #53) were selected from clusters observed at separate places whereas cluster #10 was selected as it appeared at two places.

**Figure 4:**
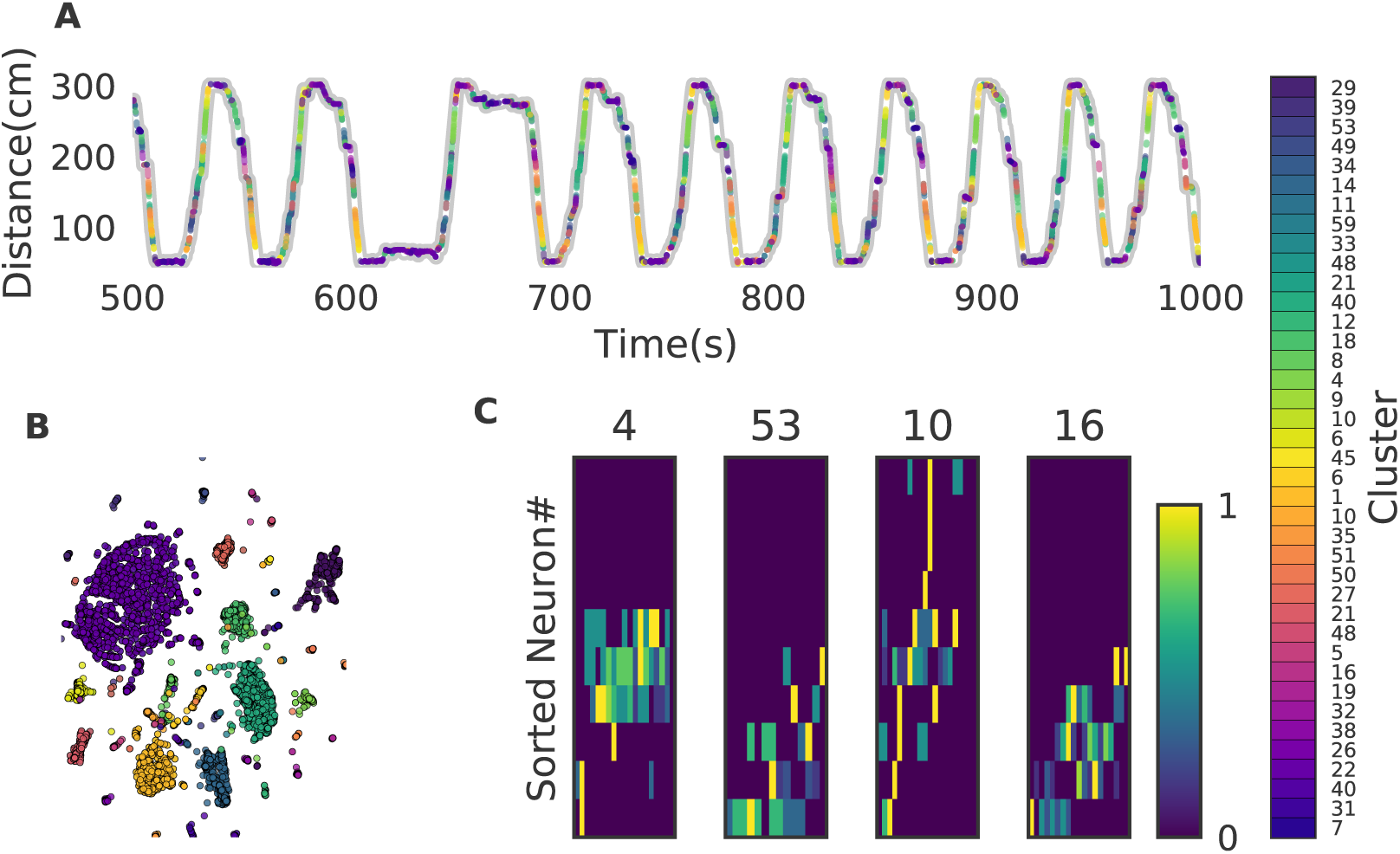
Cell assemblies extracted from hippocampal CA1. The dataset was recorded from a rat exploring a linear maze (Mizuseki et al., 2009) and is available at the data sharing site CRCNS. (A) The spatial locations in the linear maze are shown for the detected cell-assembly sequences. The x-axis shows time and y-axis shows the position of the rat. Twenty Go clusters and 20 Back clusters of data segments are shown. Each colored region shows where each segment was observed. Cluster indices are shown on the right, which were sorted and colored according the order of appearance during the going and returning along the maze. We sorted cluster indices according to their mean kernel density estimate (which is shown in Figure 6A). The window size was 100ms. (B) t-SNE visualization of the feature space. Note that most of the neighboring colors in A are also adjacent in B, suggesting that two neuronal activities observed at contiguous spatial positions have similar temporal patterns, but are still separate enough in the feature space. (C) Profiles of cell-assembly sequences are shown for each cluster. The first 10 neurons for four profiles are shown with number indicating cluster identity. Color indicates normalized firing rate of each neuron after alignment within the cluster from lowest (dark blue) to highest (yellow). Some neurons show zero rate values because their spikes were dropped from the profile during the alignment. The cells (y-axis) were sorted according to the relative temporal order (x-axis) of the peak activity of each cell in each profile. Note that the absolute length of the x-axis in each profile does not necessarily represent the actual temporal length of sequences, though the approximate length coincides the width of temporal windows (100 ms in this case). The pseudocolor code represents the firing frequency, which was normalized across neurons in the profile.

Figure 5 shows four examples of spike rasters from our extracted cell assemblies corresponding to two clusters (cluster 4 and cluster 10) together with the position and velocity of the animal. In each cluster, the spatiotemporal activity patterns vary from segment to segment, but they also resemble each other. Thus, our method is robust against changes in the temporal scale of sequences. In addition, each of the two clusters include an example of replays (at 1479 sec in cluster 4 and at 1868 sec in cluster 10) of a cell assembly in the immobile state of the animal. For the parameter values used here, these sequences were grouped into the same cluster because our method allows a certain degree of spike timing jitters. If the allowance of jitters had been narrower, they would have formed different clusters. In Figure 6A, we plotted the receptive fields of neurons and clusters when the rat was running forward, backward and stopping. It suggests that a cluster detected at a given spatial location consists of neurons that have similar receptive fields around the location. Edit similarity score for the cell-assembly sequences is significantly suppressed for shuffled neural data in which the values in each row were shuffled randomly (Figure 6B). The result suggests that the cell-assembly sequences and their profiles captured the characteristics of neural population activity.

**Figure 5:**
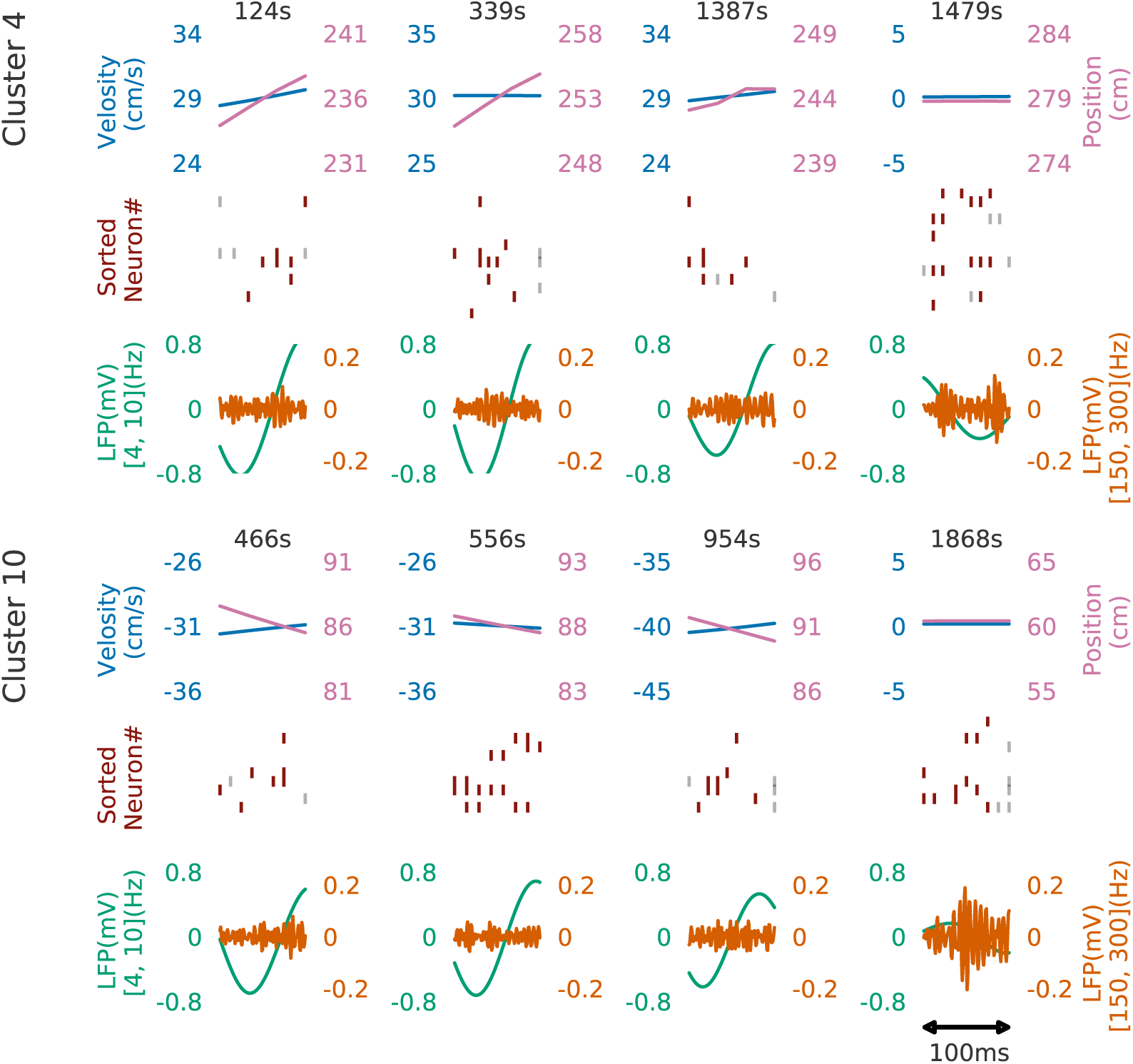
Cell assemblies detected from CA1. Spike trains are shown for four example spike sequences taken from cluster 4 (top) and cluster 10 (bottom). The top, middle and bottom panels display the velocity and spatial position of the rat, spike raster, and local field potentials band-passed at 4-10Hz (theta band) and 150-200Hz (sharp-wave ripples). In the middle panel, gray vertical bars show noisy spikes and red bars represent core spikes of the corresponding profile. Neurons are sorted according to their firing position within the average profile for each cluster.

**Figure 6:**
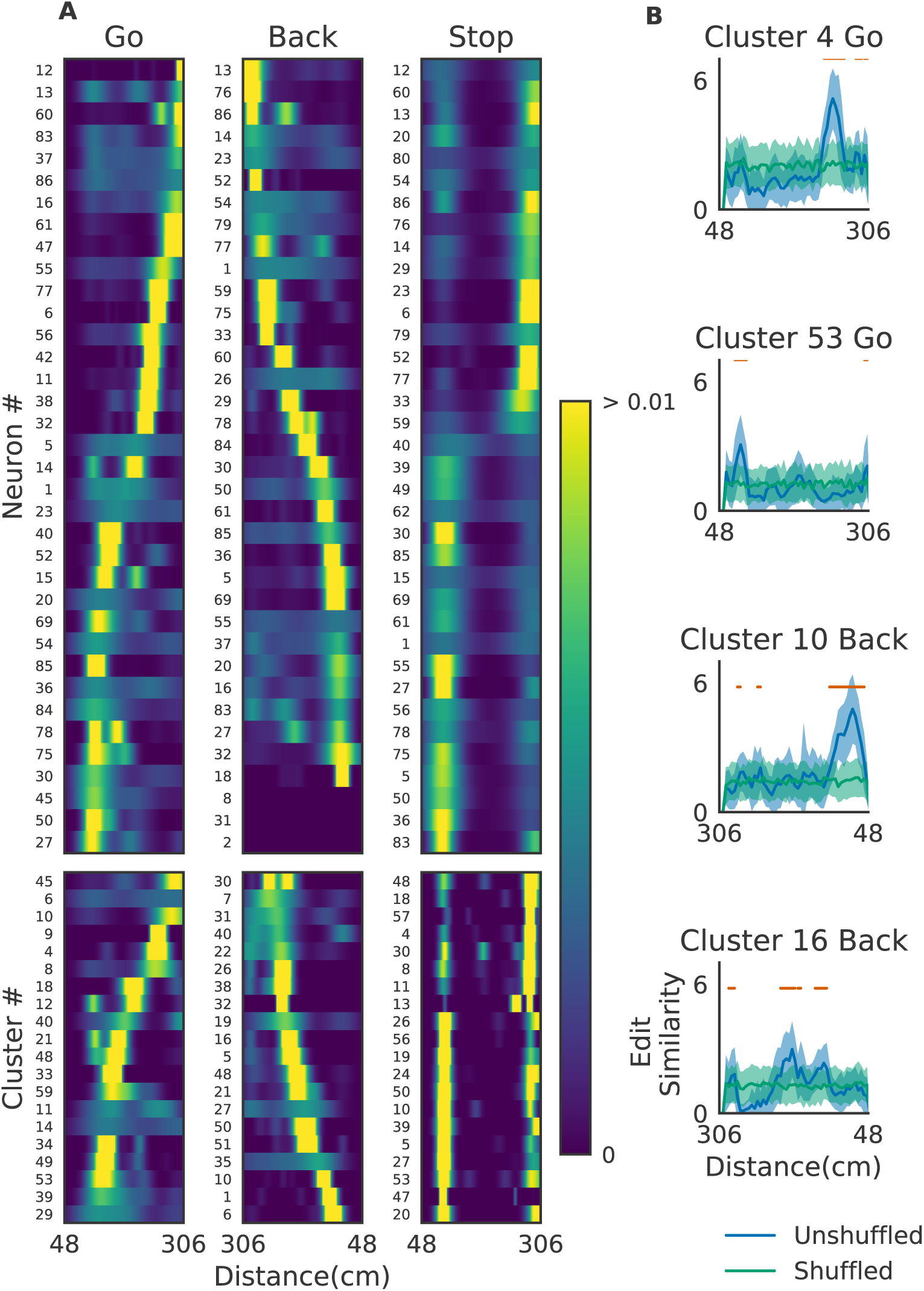
The relationships between hippocampal cell-assembly sequences. (A) The spatial receptive fields are shown for 36 hippocampal neurons (top) and cell assemblies belonging to the 20 clusters (bottom). The pseudocolor code indicates the probability of firing. (B) Edit similarity between the profiles of cell assemblies and spike trains (original and shuffled versions) of hippocampal neural population was calculated using a sliding time window.

### Cell assemblies in the prefrontal cortex

We further validated the method in neural ensemble activity recorded from the medial prefrontal cortex of rats performing a memory-guided spatial sequence task (Euston et al., 2007): see the paper for experimental details). Briefly, the rats were trained to visit eight locations equally spaced around the perimeter of a circular arena in a prescribed sequential order with electrical brain stimulation as a reward. Then, multi-neuron spike trains were recorded with a chronically implanted hyperdrive consisting of 12 tetrodes. The data contains the activity of 76 neurons and the total duration of recordings is 11,010 seconds. See Experimental Procedures for the choices of parameter values.

Our method detected 11 clusters of prefrontal cell-assembly sequences in total (Figure 7A). The previous analysis based on template matching revealed a sequence and its replay pattern in the same rat as we analyzed here (Euston et al., 2007). Though some of the detected clusters are overlapped, the larger number of detected clusters indicate that the method extracted activity patterns without any reference to events or positions on the track. These clusters were detected in both behaving state and sleep state, and some clusters were frequently replayed during sleep (Figure 7A). During the behavior, cell assemblies were typically found when the rats were approaching or leaving a reward zone (Figure 7B). The profiles of three cell assemblies are shown in Figure 7C. Each sequence usually appeared just once in a 200 ms window during behaving state, whereas they were repeated multiple times during sleep state (Figure 7D). Thus, the detected sequences were time compressed during sleep. These results are consistent with the previous findings.

**Figure 7:**
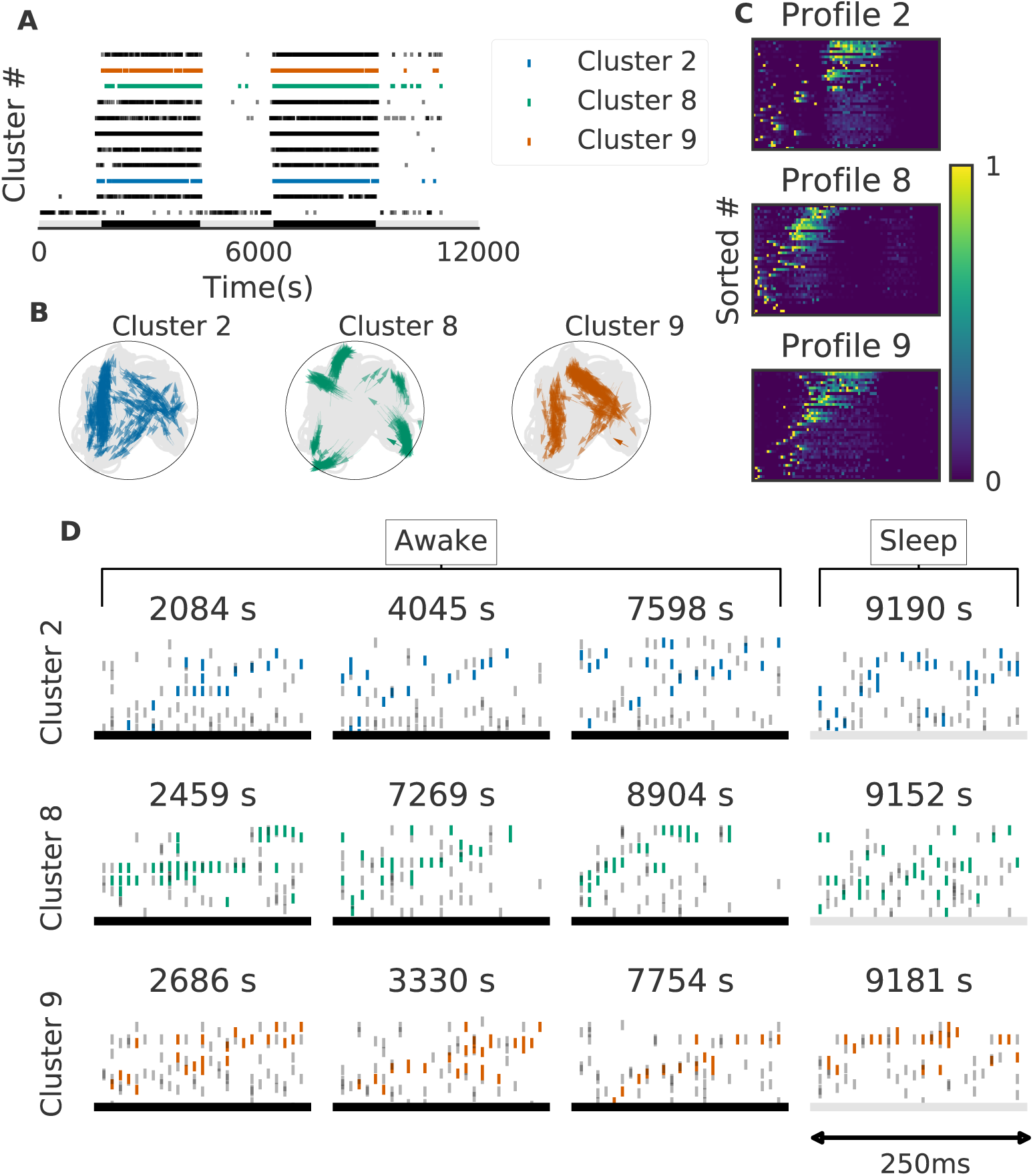
Cell assemblies detected from the prefrontal cortex. The width of time windows was 250ms. (A) The onset times of detected time windows are shown for all the clusters. (B) The spatial positions and movement directions of a rat are shown at the onset times of detected time windows belonging to three clusters. (C) Profiles are shown for three prefrontal cell assemblies in terms of the sorted neuron id and relative temporal order. The approximate length of the x-axis coincides the width of temporal windows (250 ms). (D) Cell assembly sequences detected in awake (left three panels) and sleep (rightmost panel) are shown for the three clusters. From top to bottom, each row corresponds to the profile 2, 8 and 9, respectively. Some sleep replay events showed evidence of multiple replays within the 250 ms window. This is most apparent in the first row, where the upward ramp is seen twice.

## DISCUSSION

In this study, we have developed a method for extracting multiple repeated sequences of cell assemblies from multi-neuron activity data. Our method is based on edit similarity, which was developed in computer science as a measure of similarity between strings. Edit similarity compares the serial order of common elements appearing in two strings with or without discounting variations in inter-element intervals, hence it provides a flexible and efficient metric for comparing highly noisy spatiotemporal activity patterns of cell assemblies. We have validated the method first in artificial data and then in neural activity data recorded from the hippocampus and prefrontal cortex of behaving rodents.

In the assessment with artificial data, we showed that our method is superior to PCA- (Lopes-dos-Santos et al., 2011; Peyrache et al., 2009) and ICA-based methods (Laubach et al., 1999; Lopes-dos-Santos et al., 2013). Dynamic programming (DP) based methods were previously introduced to quantify the similarity between spike trains of neurons(Victor and Purpura, 1996; Victor et al., 2007). To our knowledge, no methods have been developed with edit similarity to detect similar sequences of cell assemblies from noisy population neural data in unsupervised manner. Other methods also exist and discovered the activation of specific neuron ensembles (Chen and Wilson, 2017; Lee and Wilson, 2002; Ohki et al., 2005). The previous methods, however, are generally not effective when data has a low signal-to-noise ratio, for instance, when most of the recorded neurons do not participate in sequences. In addition, the previous methods have difficulties in distinguishing partially overlapping cell-assemblies.

In particular, our method enables blind detection of cell-assembly sequences without referring to external events such as sensory stimuli and behavioral responses. Recently, a statistical approach was proposed for the detection of cell assembly structure with multiple time scales (Russo et al., 2017). Starting from pairwise correlations in neuron pairs, the method finds significantly correlated neurons within the set of cell assemblies detected at the previous step. Acceleration of the analysis was achieved by discarding statistically less significant combinations at the next step. However, in the successive statistical tests, the detection of long sequences becomes rare and time consuming. In contrast, our method is computationally more efficient when searching longer cell-assembly sequences. It may also be inappropriate to discard long sequences just because they are statistically less significant. For instance, place-cell sequences spanning several seconds of behavior emerge in the hippocampus during spontaneous activity after spatial experience (Dragoi and Tonegawa, 2013; Grosmark and Buzsáki, 2016). We propose that behaviorally relevant cell-assembly sequences should be addressed after all possible candidates have been identified. Our method enables such an analysis of cell-assembly sequences.

It is noted that both of the examples provided here rely on highly stereotyped, repeated behaviors, which presumably entrain similar repeated patterns in the neural activity. Whether the present algorithm can be used to detect spontaneous (as opposed to stimulus- or activity-driven) patterns, such as those reported by (Luczak et al., 2007; 2009) remains to be tested. Other intriguing extensions of this method include the detection of hierarchically organized cell assemblies over multiple spatio-temporal scales. Such an extension requires a flexible on-line tuning of time windows, which is a challenge at the moment. One area where time-scaling would be particularly relevant is in the detection of replay events during sleep, which often occurs at a compressed timescale. Our method might detect considerably more reactivation events were we to adjust the temporal scaling between behavior and sleep epochs. We also note that in principle our method is applicable to optical imaging data if we can find adequate sizes of the time window and temporal discount factor.

In sum, we proposed a novel method for the blind detection of cell-assembly sequences based on the edit similarity score and an exponential discount for timing jitters. This method does not rely on the external references, hence is useful for detecting not only externally-driven firing sequences, but also internally-driven sequences emergent from arbitrary mental procedures. Whether the method reveals the involvement of cell-assembly sequences in mental processes is an interesting open question.

## ACKNOWLEDGEMENT

We thank Bruce McNaughton for his encouraging comments on the proposed method. This work was supported by Brain/Minds project by the MEXT and also partly supported by Grants-in-Aid for Scientific Research (KAKENHI) from MEXT (nos. 15H04265, 16H01289, 17H06036 to T.F.). K.W. was supported by Junior Research Associate program of RIKEN. We also thank International Neuroinformatics Coordinating Facility (INCF) for supporting this collaboration through a travel grant to M.T.

## AUTHOR CONTRIBUTIONS

T.F and K.W conceptualized and supervised the work. K.W., T.H. and T.F. developed the mathematical method and K.W. analyzed its performance. M.T. and D.E. provided the data from prefrontal cortex and contributed to the calibration of the method. T.F. and K.W. wrote the initial draft and all authors edited the manuscript.

## EXPERIMENTAL PROCEDURES

We explain three major steps of the proposed method, i.e., edit similarity score with an exponentially growing gap penalty, clustering algorithms and profile generation algorithm. The implementation of the algorithm by Python 3.6, Julia 0.5 and Bash shell script with which we carried out our analysis is available at Github (link should be added in here). We used the original implementation of the COPRA (Gregory, 2010) publicized by the author. A minor modified version of (Zhang et al., 2013) was used for the OPTICS. We have also used gnu parallel (O. Tange, 2011) for data processing.

### N-W algorithm for edit similarity

We explain the fundamentals of edit similarity score without gap penalty since this metric is not popularly used in neuroscience. Edit similarity score or edit distance quantifies the similarity between two strings with the minimum number of operations required to transform one string into the other. We can define arbitrary scoring schemes for each manipulation on strings, that is, insertion of a gap, deletion of a character, and comparison of two characters for coincidence. Needleman and Wunsch (Needleman and Wunsch, 1970) proposed one of the most widely used evaluation algorithms of this metric (N-W algorithm).

The original N-W algorithm uses DP, which essentially partitions given problem into subsequent small subproblems and composes a solution of the original problem from those of the subproblems. As an example, we evaluate the score between two strings, W(1) = ATCGTAC and *W* (2) = ATGTTAT. As shown in Figure 7, we prepare a grid (DP table) and arrange the two strings along the abscissa and ordinate of the DP table. We add a null character “#” to the heads of the two strings and fill the leftmost column and the bottom row with zeros to initialize the following iterative operation.

We assign an appropriate score to each operation (insertion, deletion and coincidence). For the sake of simplicity, in this example without gap penalty we assign + 1 to a coincidence, 0 to an insertion and a deletion. Let 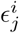 be the number of partial coincidences obtained up to the *i*-th element of *W* (1) and the *j*-th element of *W* (2). Then, we determine the value 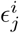 of the cell (*I,j*) of DP table by the following recursive equation:

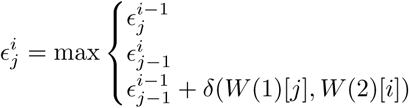

where *δ*(*i, j*) is the Kronecker’s delta: *δ* (*i, j*) *=* 1 if *i* = *j* and 0 if *i ≠ j.* We can fill the grid from the lower left cell to the top right cell with the scores calculated by the above equation (Figure 7). We note that *δ*(*x,* #)=0 for any character *x* including a null character itself. Then, we obtain the similarity score of two strings ***W***(1) and ***W***(2), which is five in this case, at the top right cell 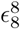. Note that the operation 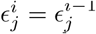 corresponds to a deletion of *W*(2)[*i*], or equivalently, a gap insertion after *W*(1)[*j*]. Likewise, 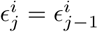 corresponds to a deletion of *W*(1)[*j*] or a gap insertion after *W*(2)[*i*], and 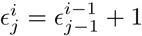 corresponds to taking a coincidence.

The DP table enables us to obtain the substring that gives the maximum number of coincidence. We can determine this substring (ATGTA in the example) by tracing arrows from the top right cell back to the bottom left cell and aligning the characters that appear after every diagonally upward move. This procedure is illustrated with gray arrows in Figure 7, and the resultant alignments of *W*(1) and *W*(2) are shown at the bottom.

### Extended N-W algorithm for neural activity

To make the N-W algorithm applicable to spike data, we make three extensions: scoring with the inner product, exponential gap penalty, and the local alignment of starting points. First, the degree of similarity between activity vectors is measured by the inner product of the vectors instead of delta function *δ*(*W*(1)[*j*], *W*(2)[*i*]) in the N-W algorithm because exact matching between two patterns is very rare in neural data. As explained in the main text, we introduced a sliding time window to get data segments of fixed length *L.* Let a matrix *W*(*t*_*k*_) be spiking activities of *N* neurons in the segment starting at time *t_k_,* and **r***_i_*(*t*_*k*_) be the column vector in the *i*-th bin of *W* (*t*_*k*_) (see Figure 1B):

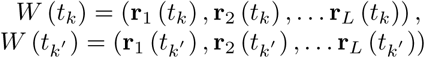

we regard *w*(*t*_*k*_) and **r***_i_*(*t*_*k*_) as a string and a character in n-w algorithm, respectively, and we evaluate similarity between activity patterns observed in two data segments at *t*_*k*_ and *t_k_’* by the inner product **r***_i_* (*t*_*k*_) *·* **r***_j_* 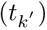.

Second, we developed a scoring scheme with exponential gap penalty, which penalizes edit similarity score with an exponential discount factor when the corresponding elements appear after different time lags in two sequences, similar in spirit to the well-known linear gap penalty scheme (Gotoh, 1982). Below, two symbols 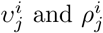 refer to the optimal numbers of vertical gap insertion and horizontal gap insertion required for partial comparison up to the *i*-th element of 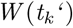 and the *j*-th element of *W*(*t*_*k*_‘), respectively. By inserting an additional gap we may earn another coincidence at the cost of an additional discount factor in the similarity. Whether one should stop or continue the insertion of a gap is determined by the comparison of the cost and benefit of the two operations. To optimize the cost-benefit balance, we should insert a maximal number of gaps that does not cost more than the benefit. After setting the initial conditions 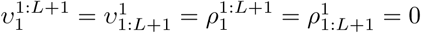 at the bottom row and the leftmost column of the table, we calculate the values of 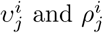 by the following recursive formula

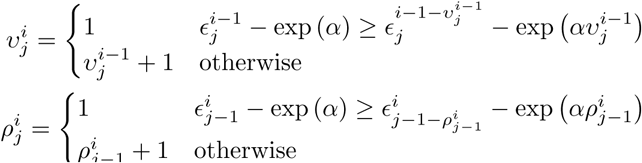

where *α* is the weight for gap penalty. The conditions

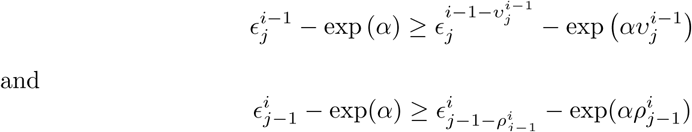

are satisfied if the cost exceeds the benefit, and then we stop insertion of a gap. The values of 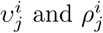 are calculated before 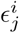 in each cell.

Third, We solved the local alignment problem by applying the algorithm proposed by (Smith and Waterman, 1981). In the case of strings (Figure 7), the heads of strings, from which we should start the comparison, are obvious. However, the heads of cell assembly sequences are not given a priori in neural data. In our scheme, when no significant coincidences are found up to cell (*i,j*) and the score in that cell is below 0, we restart recursive evaluation by setting 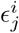 to 0. In other words, we can jump from the bottom left cell to an arbitrary cell. This scheme results in the automatic search of the optimal starting points of the comparison.

In sum, recursive equation in N-W algorithm is changed into the following rule:

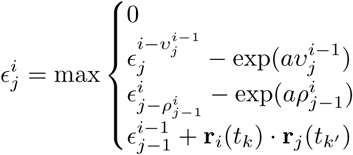

which is evaluated along with 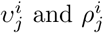. Initial conditions are given as

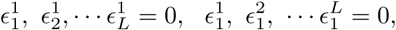

and 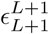 gives the maximum coincidence between the activity sequences, that is, the edit similarity score, as in the standard N-W algorithm. Below, we express 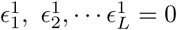 by the notation 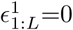.

In addition to the starting points, the end points are not given a priori, but should be determined such that the two subsequences optimally coincide with each other. We can solve this problem by backtracking of max *ε*. The procedural dependency among the cells is stored in variables 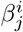: from the top to the bottom of the rules shown above, we assign “start”, “upward” (↑), “rightward” (→), and “diagonal up” 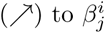, respectively. As previously mentioned, we can obtain optimal coincident substrings by back-tracking allows in the *β* table from the top right cell to the left bottom cell.

### Metric space for the clustering analysis

The algorithm described above gives a similarity matrix consists of edit similarity scores *E*(*k,k*’) between pairs of data segments *W*(*t*_*k*_) and *W*(*t*_*k*_’). We calculate a distance matrix as *D*(*k,k’*) = max(*E*(*k,k*’))-*E*(*k,k*’), where the maximum is taken over all possible pairs of segments. The distance matrix defines a distance metric in a high-dimensional feature space in which data segments containing similar activity patterns are distributed at neighboring locations. As demonstrated in Figure 2, we can extract similar cell-assembly sequences through the clustering of data points in this feature space.

### Dimensionality reduction by t-SNE

t-Distributed Stochastic Neighbor Embedding (t-SNE) is an algorithm that maps high-dimensional data into a low dimensional space, typically two or three-dimensional space while maintaining the original data structure in the high-dimensional manifold. It accomplishes the mapping by reducing the Kullback-Leibler divergence between *p*_*ij*_ and 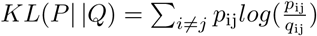, where *p*_*ij*_ is the conditional probability distribution between two data points labeled by *i* and *j* in the original manifold and *q*_ij_ is the conditional probability distribution of the corresponding data points in the embedded manifold. This algorithm uses a normal distribution for *p*_ij_ and a Cauchy distribution for *q*_ij_ to faithfully preserve the data distribution in the original space.

### Tricks for reduction of the computational cost

Evaluation of the similarity matrix is generally quite heavy; the complexity is O(M^2^) where *M* is the total number of data segments. To avoid the heavy computation, we reduced the number of the matrix elements to be evaluated by an algorithm proposed by (Cohen et al., 2001). In this procedure, we reduce the calculation of edit similarity for pairs of data segments that do not share active neurons. Let *w* be a Boolean matrix in which the element (*i,k*) is 1 if neuron *i* fires at least once in the segment *W*(*t*_*k*_) or otherwise 0:

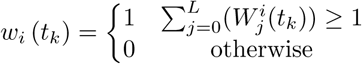

Our goal is finding pairs of data segments yielding similar column vectors *w*(*t*_*k*_) and *w*(*t*_*k*’_) with low computational cost. We measure the similarity between *w*(*t*_*k*_) and *w*(*t*_*k*’_) by Jaccard similarity defined as

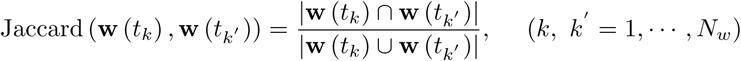

where |x∩y| denotes inner product of given two vectors, |x∪y| counts number of non-zero element of the sum of them. The value of Jaccard similarity is between 0 and 1, and is close to unity if the column vectors at time *t*_*k*_ and *t*_*k*’_ are similar.

Because calculation of Jaccard similarity for every possible pair of vectors is also O(M^2^), we wish to find out pairs that are likely to give highly similar *w*(*t*_*k*_) without direct calculation. For this purpose, we can make use of the statistical properties of Jaccard similarity. Now a trick is to use hash function 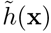 which randomly assign an integer to the given integer/vector x. Throughout this study, we used a built-in hash function of programming language Julia. We define function 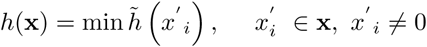 that returns the minimum hashed number made by non-zero elements of **x**. The value is called the minimum hash (min-hash) value. Importantly, we can prove the following relationship (Cohen et al., 2001):

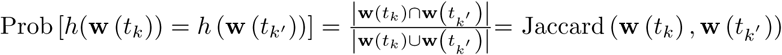

With this relationship, we can obtain Jaccard similarity without pair-wise comparisons of column vectors:

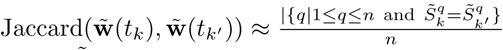

where 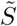 is called a signature matrix that contains the min-hash values in different hash functions, i.e., 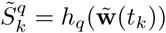, and *h*_*q*_ is the *q*-th hash function and *n* is the number of hash functions. In this matrix, elements in a column are min-hash values of a data segment generated with different hash functions, and elements in a row are min-hash values of all data segments generated with a hash function.

To further reduce computation, we used the banding technique in the evaluation of Jaccard similarity (Cohen et al., 2001). We divided 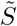 into *b* bands of *l* rows each, thus *n=bl.* Suppose that two vectors 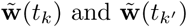 have Jaccard similarity *s*, then the probability that the min-hash signatures of two columns coincide at least in one row of the matrix is *s*. Then, the probability that the signatures of two columns are identical in all rows of at least one band is 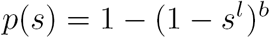, which is an *S*-shaped function of *s* and hence can be used for determining a threshold value of the similarity. We hash all the bands, and search bands in which two columns have the same hash value. The only pairs of data segments that have the same hash value in more than one band are used for similarity matrix calculation. For instance, *p*(0) =0.000, *p*(0.3) =0.007, *p*(0.5)=0.091, *p*(0.7) =0. 424, *p*(0.8) =0.696, *p*(0.9) =0.931, and *p*(1.0) =1.000 when *b =* 5 and *l* = 2. In the present analysis, the values of *b* and *l* were dynamically adjusted by data itself. We explain the method for the adjustment in the next section.

### A policy for division of signature matrix

To apply the above algorithm to neural data, we employed a heuristic method to determine two parameters for Jaccard similarity (*b* and *l*) from neural data. We calculated the average firing rates of individual neurons over the entire length of data, and we determined *b* and *l* assuming independent Poisson spiking neurons having the same firing rates. The parameter for a smaller threshold (Jaccard1) gives the similarity expected under the assumption of independent Poisson spiking, whereas a parameter for a larger threshold (Jaccard_2_) represents the similarity expected when the two data segments contain sequences with a certain length. Let #*_i_* be the total number of spikes of neuron *i* during the interval [0, *T*]. From #*_i_*, we can calculate the probability that neuron *i* has at least one spike in the segment 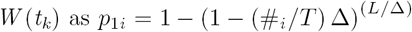, where ∆ is the size of a bin. Then, the index 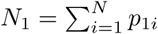 is the expected number of active neurons within the time window. Then, the expected number of coincidently active neurons in an arbitrary pair of data segments is 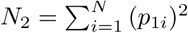, and Jaccard_1_ is calculated as N_2_/(2*N*_1_-*N*_2_). Now, suppose that two data segments contain additional *N*_3_ coincidently active neurons. In this case, the expected Jaccard similarity, or Jaccard_2_, is given as (*N*_2_+*N*_3_)/(2*N*_1_−*N*_2_+*N*_3_). In this study, we searched such values of *b* and *l* that keep the probability 1 − (1 − *s^r^*)*^b^* sufficiently high compared with Jaccard_1_ and sufficiently low compared with Jaccard_2_.

The proposed method is different from the standard approach based on statistical analysis and remains somewhat heuristic. However, the aim of this method is to reduce the load of heavy computation for large neural data without losing candidate sequences, as we consider that the behavioral importance of detected sequences should be analyzed based on their relationships to behavioral data.

### Construction of profiles for clustered sequences

Here, we explain our iterative multiple alignment algorithm for constructing profiles of clusters. It is based on the algorithm by (Barton and Sternberg, 1987). In the original algorithm, we initialize the algorithm with a tentative profile, which is obtained by taking the longest common subsequence between the two data segments in a cluster that show the highest match in edit similarity. After the initialization, we search a next data segment that gives the most similar profile to the tentative one, and update the tentative profile using edit similarity. We repeat this procedure until the tentative profile converges.

In our method, we made two major modifications to the original algorithm. First, we chose two arbitrary data segments in the initiation step to reduce the computational cost. The final results did not significantly differ between our approach and the original one. Second, in generating a profile, we used the z-score of spike count in each data segment. Namely, for each neuron we calculated the average and variance of spike count per bin over the data segment, and then subtracted the average from spike count in each bin and normalized the difference by the variance. The use of z-score suppresses the influences of highly active neurons on the detection of ensemble firing sequences. Finally, in each step, a Gaussian filter with mean 0 and variance *σ* was applied to the tentative profile. Variance *σ* was initially as large as the window size and gradually reduced to the bin size as iterations proceeded. This filtering prevented a profile from containing more than two similar sequences, thus enabled a robust detection of minimal sequences.

### F-score for artificial data analysis

Purity and Inverse Purity are defined as

Purity = 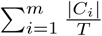 Precision,

Inverse Purity = 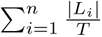 Precision,

where *m* is the number of detected clusters *C* = {*C*_1_, *C*_2_, …, *C*_*m*_}, *n* (=11) is the number of true clusters and a noise cluster in the artificial data *L =* {*L*_1_, *L*_2_, …, *L*_*n*_}, and *T* is the total number of data points (i.e., segments of spiking data). The noise cluster consists of spurious cell assemblies. Precision(*C*_*i*_, *L*_*j*_) is defined as 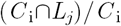, which represents the fraction of members of the *j*-th true cluster in the *i*-th detected cluster. In the above expressions, weights are determined such that a larger cluster contributes more strongly to the weighted sums. We note that Purity and Inverse Purity take their values within the interval [0, 1]. F-score is defined as the harmonic mean of Purity and Inverse Purity to penalize two trivial solutions. In one such solution, each data point constitutes an independent cluster (i.e., *m = T*). In the other solution, all data points are classified into a large cluster. In these trivial cases, Purity, but not Inverse Purity, takes the maximum value of unity.

In the evaluation of our method with simulated data, we searched the optimal clustering that maximizes the F-score by gradually changing threshold for the agglomeration/division of data points. We optimally separated the target cell assemblies from spurious cell assemblies. In evaluating the PCA- and ICA-based methods, we calculated overlaps between the population activity vector and the principal (or independent) components in all data segments. Then, we searched for a segment that showed the highest value of the overlaps, and the component yielding the highest value was associated with the cell assembly that was activated in the segment. The detected data segments were removed from the next search. The above search procedure was repeated until all cell assemblies were associated with some components without overlapping. The remaining components had no partner cell assemblies and were regarded as noise cluster. Thus, we obtained *n =* 11 clusters including a noise cluster of spurious cell assemblies.

### Parameter choices

Here we list the values of parameters in our algorithm. For the artificial data, we have tested multiple parameter searched different parameter settings to find the values that record the maximum performance. Each parameter tested within the following range; the power of exponential growing gap penalty *α*=1.0; the number of points for a minimal cluster in OPTICS *MinPts 2* to 20 (Ankerst et al., 1999); parameter for COPRA *v 2* to 20(Gregory, 2010).

For the hippocampal data, the following parameter values were used: the power of exponentially growing gap penalty *a* = 0.1; the length of sliding time window *T_w_=* 100 (ms), which is the typical period of one cycle of theta oscillation; criteria for fast computation *minlen =* 10; the number of points for a minimal cluster in OPTICS *MinPts =* 20; parameter for COPRA *v =* 4.

The prefrontal data were analyzed with various widths of time windows ranging from 250 ms to 2.5 sec because the characteristic time scale of sequences were not known. All the results shown here are obtained for the width of 250 ms, which yielded reasonable sequences. The temporal discount factor was set as *a* = 0.03. The number of points for a minimal cluster in OPTICS *MinPts =* 400; parameter for COPRA *v =* 30.

